# Mapping Hsp104 structure and substrate interactions using crosslinking mass spectrometry

**DOI:** 10.1101/2022.08.19.504539

**Authors:** Kinga Westphal, Lukasz A. Joachimiak

## Abstract

Molecular machines from the AAA+ family play important roles in protein folding, disaggregation and DNA processing. Recent cryo-EM structures of AAA+ molecular machines have uncovered nuanced changes in conformation that underlie their specialized functions. Furthermore, complexes between these machines and substrates begin to explain their mechanism of activity. Here we explore how crosslinking mass spectrometry (XL-MS) can be used to interpret changes in conformation induced by ATP and how substrates are associated. We applied a panel of crosslinking reagents to produce high-resolution crosslinking maps and interpret our data on previously determined X-ray and cryo-EM structures of Hsp104 from a thermophilic yeast, *Calcarisporiella thermophila*. We developed an analysis pipeline to differentiate between intra-subunit and inter-subunit contacts within the hexameric homo-oligomer. We identify crosslinks that break the asymmetry that are only present when ATP is bound and are absent in an ATP-binding deficient mutant. Finally, we identify contacts between Hsp104 and a model substrate to identify contacts on the central channel of Hsp104 across the length of the substrate indicating that we have trapped interactions consistent with translocation of the substrate. Our simple and robust XL-MS-based experiments and methods help interpret how these molecular machines change conformation and bind to substrates even in the context of homo-oligomeric assemblies.

## Introduction

In living organisms, molecular chaperones play important roles in folding, refolding and prevention of protein aggregation. Molecular chaperones, by interaction with their client substrates prevent their aggregation, enable them to fold properly thus retaining their functional shape.[1] Yeast heat shock protein 104 (Hsp104) along with its bacterial homolog caseinolytic peptidase B (ClpB) belongs to a AAA+ (ATPases Associated with diverse cellular Activity) superfamily of protein disaggregases that employ the energy from ATP hydrolysis to disassemble and remodel misfolded proteins.[2] Such molecular chaperones are especially important in maintaining the protein homeostasis, disturbance of which may cause several protein neurodegenerative diseases such as Alzheimer’s, Parkinson’s and Huntington’s.[3] In yeast, Hsp104 cooperates with Hsp70 and Hsp40 to maintain its disaggregating activity.[4] Hsp104 in yeast has been implicated in disassembling prions including sup35 which can promote prion propagation.[5] Under normal conditions Hsp104/ClpB concentration is rather low and significantly increases when stressful stimuli occur, which induces its ability to mediate misfolded protein disaggregation.[6] In contrast to bacteria, fungi and plants, animal cells do not encode a classical Hsp104 orthologue, however recent data indicate that a AAA+ protein valosin contain protein (VCP) or p97 may be implicated in disaggregation of tau and alpha-synuclein aggregates to promote seed formation in disease.[7] Thus, mechanistic knowledge on these molecular machines may yield exciting insight into control of protein disaggregation with direct implications in understanding neurodegenerative diseases.

Like other AAA+ proteins, Hsp104/ClpB is a two-tiered ring-shaped hexamer with an axial channel, which plays a key role in ATP-driven translocation and renaturation of aggregates.[8] All molecular chaperones are divided into subfamilies based on their architecture and function (Hsp40, Hsp60, Hsp70, Hsp90, Hsp100 and small Hsp).[9] Hsp104 is assembled into a homo-hexameric complex comprised of 102 kDa subunits that consists of 908 residues belonging to the Hsp100 family.[6] Both Hsp104 and ClpB are composed of the following domains: (i) N-terminal domain (NTD)[10] (cooperates in substrate translocation), (ii) two nucleotide-binding domains (NBD1 and NBD2),(interact with substrate and drive translocation), (iii) middle domain (MD) (indispensable in substrate disaggregation and interaction with Hsp70), and (iv) C-terminal domain (CTD) (typical only for Hsp104, that is essential for oligomer assembly). Fig. 1a [11,12] Within Hsp100, two classes of proteins can be distinguished: (i) class 1 – proteins that contains two Walker-type nucleotide binding domains (NBDs), like Hsp104 and ClpB and (ii) class II which includes these proteins that only have one NBD domain. It is believed that these two NBDs domains coordinate the binding and translocation of the misfolded protein via central channel of Hsp104 hexamer.[13] They contain highly conserved tyrosine residues, which shove substrate through the channel in hydrolysis-driven manner.[11] Such a mode of action induces conformational changes of the particular domains that taking part in translocation and variables in the channel diameter. [10,14] As a consequence, active high-resolution hexameric form of Hsp104 remains difficult to capture, which renders understanding the mechanism of its action. Recently a series of cryo-EM structures have uncovered asymmetry in the assemblies revealing a “staircase” arrangement and putative mechanism of substrate binding and threading through the central pore. [11,15-18]. Moreover, combined crystallographic and cryo-EM studies on the same Hsp104 complex from *Calcarisporiella thermophila* (herein ctHsp104) revealed more nuanced conformational changes. [19] It remains unknown how these molecular machines recognize substrates to promote the initial binding that leads to subsequence unfolding and threading of the unfolded chain through the central pore.

**Figure 1.**
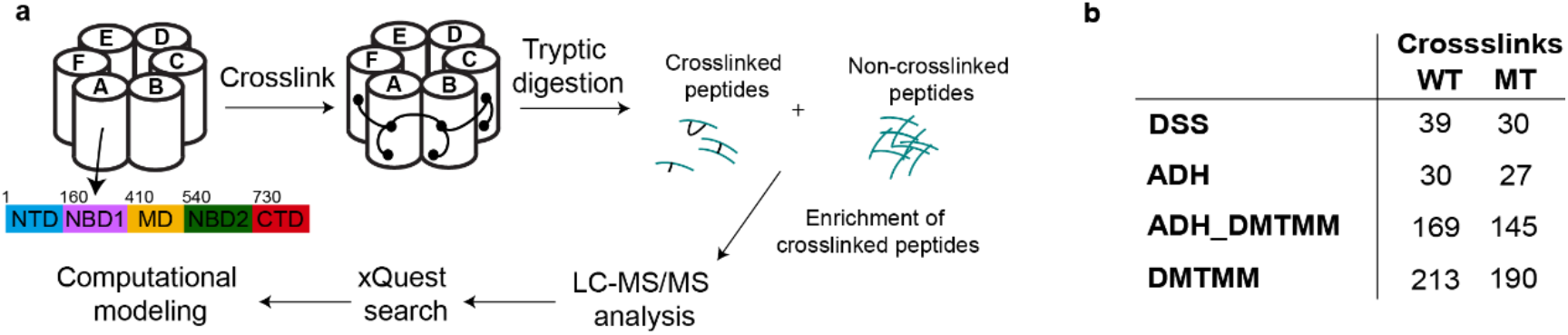
Mass spectrometry analysis of cross-linked Hsp104. **a**. Both Hsp104 wild type (wt) and Hsp104 mutant (mt) were cross-linked using the following crosslinking agents: (i) DSS; (ii) ADH and DMTMM; (iii) DMTMM. For that purpose, Hsp104 wt/mt was treated with the particular crosslinking reagent and enzymatically digested to yield a pool of crosslinked and non-crosslinked peptides. All cross-linked peptides were chromatographically enriched and analysed using LC-MS/MS. The identification of the peptides and linked (affected) chains was performed with the use of xQuest.[42] **b**. The number of all crosslinks formed in Hsp104 wt/mt after treatment with particular crosslinking agent.

Here we took advantage of ctHsp104 which has been resolved using both X-ray crystallography and cryo-EM and applied a combinatorial crosslinking mass spectrometry approach to validate our crosslinking data. Additionally, we use these data to interpret changes in conformation due to ATP binding/hydrolysis. Using the identified crosslinks from three different chemistries on Hsp104 with ATP or an active site mutant Hsp104: (1) DSS, (2) DMTMM/ADH and (3) DMTMM and employing a novel analysis pipeline to test different spatial crosslink geometries we show how the crosslinks can be used to uncover symmetric and asymmetric interactions. Finally, we employ our XL-MS approach to map how a model substrate, PCSK9, binds to Hsp104 and identify contacts points that track how the substrate traverses the pore. Our data supports that our novel crosslinking approach and pipeline can be used to interpret conformational changes in the molecular machines in their interactions with substrates or ligands.

## Results

### Strategy to interpret XL-MS data on experimentally determined structures of Hsp104

The molecular chaperone Hsp104 is comprised of six identical subunits arranged in a ring. To gain molecular insight into the structure and asymmetry of the assembly we employed a high-resolution crosslinking mass spectrometry approach (Fig. 1a) leveraging several different chemistries including disuccinimidyl suberate, adipic dihydrazide (ADH)/4-(4,6-dimethoxy-1,3,5-triazin-2-yl)-4-methyl-morpholinium chloride (DMTMM) and DMTMM alone on wildtype (ctHsp104 wt) and 383P→A mutant unable to bind ATP (herein ctHsp104 mt). Proline at residue 383 interacts with adenine and ribose of ATP molecule. [19] We identified 451 and 392 crosslinks across the different chemistries for wt and mt ctHsp104 (Fig. 1b). For the Hsp104 from this species, there are two structures available that were determined by X-ray crystallography and cryo-EM and revealed a left-handed helical assembly with unprecedented atomic detail [19]. We next decided to interpret our large crosslinking dataset using these two experimentally determined structural models. While these Hsp104 assemblies are homo-oligomeric there are three possible crosslink geometries between a pair of residues: intra-subunit (intramolecular; I), inter-subunit forward (IF) and inter-subunit reverse (IR) (Fig. 2a) when looking at three contiguous subunits (Fig 2a, F, A and B). Furthermore, if we assume any asymmetry between the subunits, we must consider 6 possible intra, inter-forward and inter-reverse geometries.

**Figure 2.**
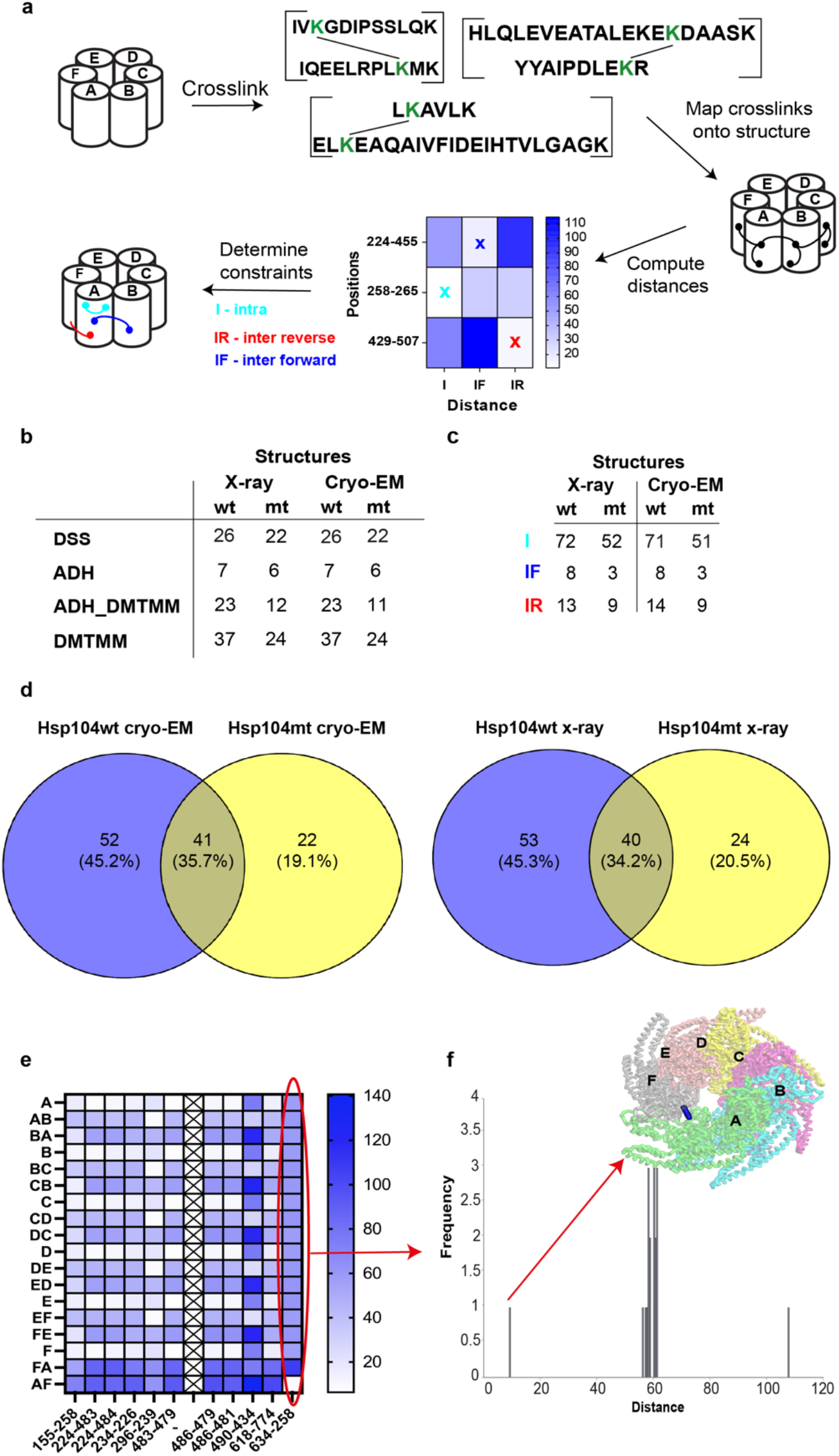
Analysis of mass spectrometry derived crosslinks on experimental structures of Hsp104. **a**. With the use of particular crosslinking agent, we determined the restraints (e.g., formed between lysine residues). Using xwalk [43] all distance restraints were computed and mapped onto Hsp104 X-ray and cryo-EM structure. The main purpose was to determine the type of crosslink, whether it is intra-crosslink (I) (formed within one subunit) or inter-crosslink (formed between two neighbouring subunits). Moreover, it enabled to establish the subtype of inter-crosslink, whether it was formed clockwise (inter forward (IF)) or counter-clockwise (inter reverse (IR)). All distance values were grouped using heat maps – for each restraint three distance values corresponding to I, IF and IR crosslinks were included. The brighter the colour, the shorter the distance between two linked residues. In each case, the shortest distance value determined the type of crosslink, which might be I, IF or IR, respectively. The other cross-links, which distance values exceeded the admissible value were excluded from cross-linking data. **b**. Summary of crosslinks formed in Hsp104 wt/mt after treatment with particular crosslinking agent, which distance values are acceptable. The results revealed the similarities between X-ray and cryo-EM structures. As it can be noticed, the number of cross-links formed onto Hsp104wt cryo-EM and X-ray structure as well as Hsp104mt cryo-EM and X-ray structure is comparable. **c** Summary of crosslink types formed in Hsp104 wt/mt after treatment with particular crosslinking agent. These data also confirm that cryo-EM and X-ray structures are similar. The number of sub-types of cross-links formed onto Hsp104wt cryo-EM and X-ray structure as well as Hsp104mt cryo-EM and X-ray is almost the same. **d**. Overlap of all the crosslinks identified in each of the two (Hsp104wt and Hsp104mt) datasets. **e**. With the use of heat maps, it was possible to easily find distinctive cross-links. In this case, only crosslink formed between 634 and 258 residues of A and F units of Hsp104 fits perfectly the structure. **f**. The 634-258 constraint is the only crosslink, which distance value does not exceed 20 A, while the majority of restraints formed between or within other subunits do not fit onto the Hsp104 structure. This crosslink, formed between A and F subunit was mapped onto Hsp104 structure as a representative of distinctive restraint. Each of the protein units has its own colour to facilitate identification of particular parts of the Hsp104 hexamer.

### Mapping identified crosslinks onto discrete subunit geometries

We set out to interpret our comprehensive XL-MS dataset derived from three different crosslinking chemistries on wt and mt ctHsp104 (Fig. 1a) in the context of two experimental structures determined by X-ray crystallography and cryo-EM [19]. All the crosslinks were also mapped onto all possible combinations of geometries for a single subunit (I, i.e. A) and two geometries between subunits, inter-subunit forward (IF, i.e., AB) and inter-subunit reverse (IR, i.e. BA). A subset of the identified crosslinks 213 of 451 (ctHsp104 wt) and 190 of 392 (ctHsp104 mt) can be mapped on resolved parts of the experimental structures. This calculation was performed for all single subunits and all local subunit pairs (A, AB, BA, B, BC and CB, etc…). We first ascertained how well our identified crosslinked pairs fit on each structure by calculating the C_alpha_-C_alpha_ distance for all possible subunit geometries (Fig. 1b, e.g. I, IF and IR) initially focusing on the X-ray structure. Since each residue can form a crosslink with a second residue within a subunit (I) or neighboring subunit (IR or IF) it was essential to first classify all crosslinks and identify the most probable geometry based on distance (for details see Fig. 2a). To simplify interpretation of this data we used heat maps to order all the values (see Supplementary Fig 3-6), thus enabling precise assignment of contacts based on subunit and crosslinker geometry to those shorter than 30 Å (for DSS) and 15 Å (for ADH/DMTMM). All identified crosslinks could be mapped onto both X-ray and cryo-EM structures, despite the fact that in these two modeled structures of ctHsp104 the part of C-terminal sequence was missing (882 out of 908 aa). As a result, we determined geometric assignments for all the probable crosslinks (93) with specific geometries that mapped onto modeled parts of the X-ray structure of ctHsp104 from the ctHsp104 wt XL-MS cumulative datasets and similarly, we were able to assign 64 contacts to specific geometries from the ctHsp104 mt. The numbers of explainable crosslinks were similar when mapped onto the cryo-EM structure (Fig. 2b). These values are consistent with our calculated false discovery rates (Supplementary Fig. 2). Of all the crosslinking chemistries tested, the most abundant crosslinks were formed using DMTMM, which traps zero-length crosslinks between lysine residues to carboxylic groups of glutamic or aspartic acid for which we identified 37 and 24 crosslinks that are mappable onto the X-ray structure (see Fig. 1b). From our calculations we also partitioned the contacts into I, IR and IF to determine which contacts are the most frequent in our dataset. The data revealed that intramolecular crosslinks within a subunit constitute the most common type of linker, in both cases intra crosslinks represent around 80% of all constraints formed in Hsp104 wt and Hsp104 mt. The rest of the crosslinks were formed between two neighboring units (see Fig. 2c). We also find that the majority of intramolecular crosslinks localized to MD and NBD1 domains as well as the inter-subunit contacts (see Fig. 3) It is perhaps not surprising that intramolecular contacts are the most abundant contacts as these should be the least dynamic. Partitioning of the IR and IF inter-subunit contacts is less interesting because it is purely derived from the ordering of the pairs from the rather arbitrary assignment of the two peptides. The assignment of the first (A) and second (B) peptides in the software where peptideA-peptideB contacts can be attributed to the IR inter-subunit but are equivalent to peptideB-peptideA in the IF inter-subunit arrangement. Together, our data and analysis suggest that we can evaluate our crosslinking data in the context of experimental structures even when applied to homo-oligomeric assemblies enabling assignment of specific subunit geometries. All the other crosslinks that fit onto Hsp104 structures are presented in Supplementary Figure 9.

**Figure 3.**
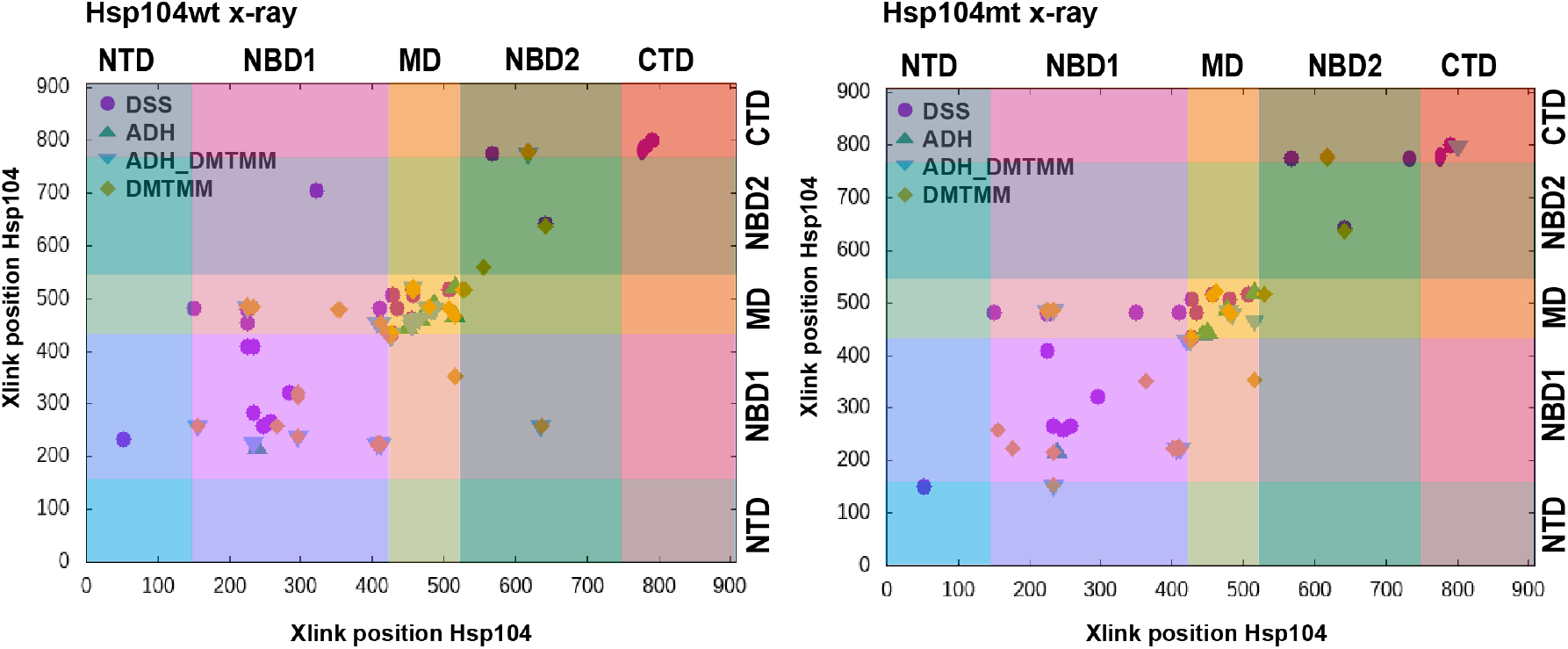
Hsp104 regions that predominantly form crosslinks. All crosslink positions were plotted against each other to reveal Hsp104wt/mt regions that are involved in the formation of constraints. In each case, the highest number of crosslinks was formed within NBD1 and MD domains. Only minor differences can be detected between genetic variations of Hsp104 (Hsp104mt forms fewer constraints), while crosslink distribution mapped onto particular structures does not reveal significant differences, which confirms structural similarity of Hsp104 cryo-EM and X-ray structure. For cryo-EM variants see Supplementary Figure 8.

### Identification of XL-MS contacts consistent with asymmetric subunit geometry

It is also worth noting that the XL-MS datasets derived from the Hsp104wt samples yielded a greater number of crosslinks compared to Hsp104mt, which may suggest that the Hsp104wt structure is less dynamic because it is able to bind ATP which can facilitate trapping crosslink contacts while the ctHsp104 mt is unable to bind to ATP. Indeed, changes in dynamicism in response to ATP binding have been previously noted in other molecular machines such as the eukaryotic chaperonin TRiC/CCT (ref).[20] In order to discover other differences between wt and mt ctHsp104 based on XL-MS patterns, we compared overlap of all the crosslinks identified in each of the two datasets and find that 40 crosslinks are the same across the two datasets with only 53 and 24 different for Hsp104 wt X-ray and Hsp104 mt x-ray, respectively (Fig 2d.) Similar values were obtained for Hsp104 wt/mt cryo-EM – 41 crosslinks are the same, while 52 and 22 are different, respectively. Among all the constraints, the most distinctive crosslink that was uniquely identified in the wt Hsp104 dataset was consistent with the geometry between subunits FA linking residues 634 and 258 on the X-ray structure of ctHsp104. Interestingly, this contact is incompatible with the other subunit geometries (i.e. AB, BC, CD, DE, EF) where the distances are larger but because the helical twist begins at subunit A and ends on subunit F (i.e. defined by the “staircase” arrangement of the hexamer) the spacing between these two subunits is smaller (see Fig. 2 e,f). Furthermore, this crosslink distance was consistent with the crosslinker geometry in both the cryo-EM and X-ray structures, which confirms that these two structures, although determined by different techniques, are in fact similar and consistent with our XL-MS analysis of Hsp104 in solution. Because this crosslink was not identified in the mt Hsp104 crosslink dataset, it could indicate that the ATP-binding deficient mutant may not be able to sample these FA contacts as readily. Thus, our data suggest that we have been able to identify a unique interaction that is sampled more frequently (i.e., stably) in the wt Hsp104 complex but less frequent in the mt Hsp104. We suspect that this is largely driven by ATP binding activity which may rigidify the assembly while the ATP-binding deficient mutant is more dynamic.

To highlight how a contact fits one specific subunit geometry over others we illustrate this visually by mapping the distances on a subunit pair (i.e., AB) (Fig. 4). The crosslinks for which distances are longer than the permissible distance (based on geometry of the crosslinker) were marked with red stick while the crosslinks below the acceptable distance threshold were marked with a black stick. In 4a case, 224-479 crosslink mapped in an intra (B-B), inter forward (B-C) and inter reverse (C-B) combinations confirmed that the constraint distance value is only acceptable for interaction within B unit – distances of B-C and C-B crosslinks exceeded the assumed limit value. For the 224-410 crosslink, only B-C linkage was acceptable (Fig. 4b), while the crosslink 410-481 represents the formation of inter reverse constraint (C-B) (Fig. 4c).

**Figure 4.**
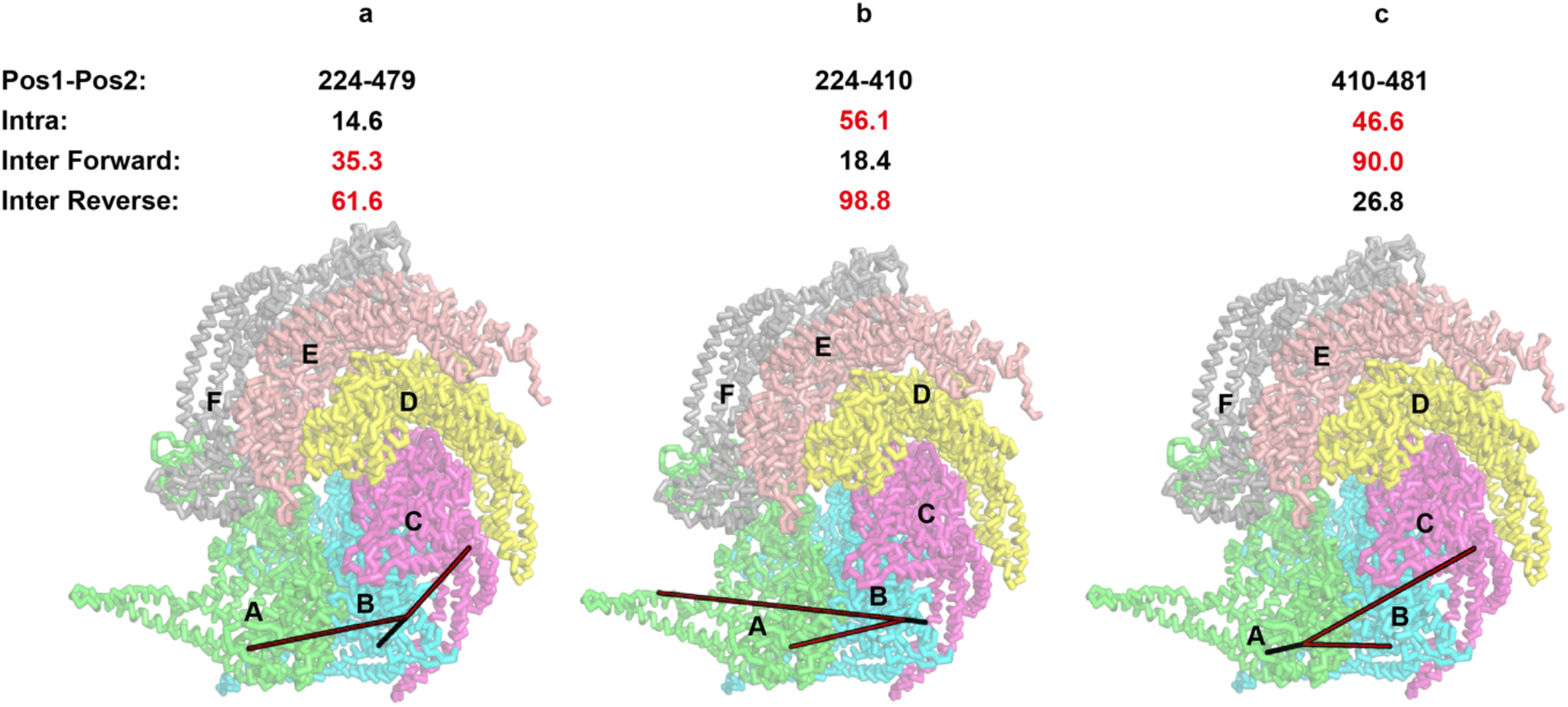
Graphical representation of strategy to discriminate geometrically compatible crosslink. The selected crosslink was mapped onto Hsp104 structure as a representative of each type of constraint. The crosslink was mapped onto the structure in each configuration: **a**. within B unit, which corresponds to intra (I) crosslink, **b**. between B and C units, which represents the intra forward (IF) linkage and **c**. between B and A units, which depicts the inter reverse (IR) constraint. On the left side, the presented crosslink formed acceptable linkage only within B unit (black lines), while BC and BA distances were too long and unacceptable (red lines). In the exact same way, the middle structure and the right structure present the formation of inter forward (IF) and inter reverse (IR) cross-link, respectively.

### Probing Hsp104-substrate interactions by XL-MS

The recent cryo-EM revolution has yielded an explosion of structures of molecular machines [21-28]. For many years these machines, including Hsp104, were modeled using symmetric averaging but new developments in instrumentation and methods have allowed reconstruction of asymmetric structures in different conformations. [11,29] Additionally, these structures have begun to uncover how substrates may bind to Hsp104 and thread through the pore. [18] Here we attempted to leverage high-resolution XL-MS to trap intermediates of a model substrate, PCSK9, bound to ctHsp104. Hsp104 was recombinantly co-expressed with PCSK9 and purified using a standard protocol. The ternary complex was reacted with DSS, DMTMM/ADH and DMTMM and the samples were processed using our XL-MS pipeline. We identified 24 intermolecular crosslinks formed between ctHsp104wt and PCSK9 with low FDRs (Fig. 5b). The majority of crosslinks identified were derived from the DMTMM chemistry (Fig. 5c, green triangle) and moreover almost each region (i.e., domain) of Hsp104 formed crosslinks with PCSK9. To reveal which Hsp104 residues are precisely involved in the interaction with the model substrate, we determined short regions, comprising of around 100 amino acids, that bind to different residues of PCSK9. In the Hsp104 sequence, residues that predominately link with PCSK9 are located within two ranges: (i) 100-200 aa and (ii) 400-500 aa (Fig. 5d) that correspond to the NTD and MD domains. This is consistent with initial recruitment of substrate to surfaces on the outside of Hsp104. Indeed, these domains have been implicated in binding to substrates using biochemical approaches as well as structural approaches. [30,31]

**Figure 5.**
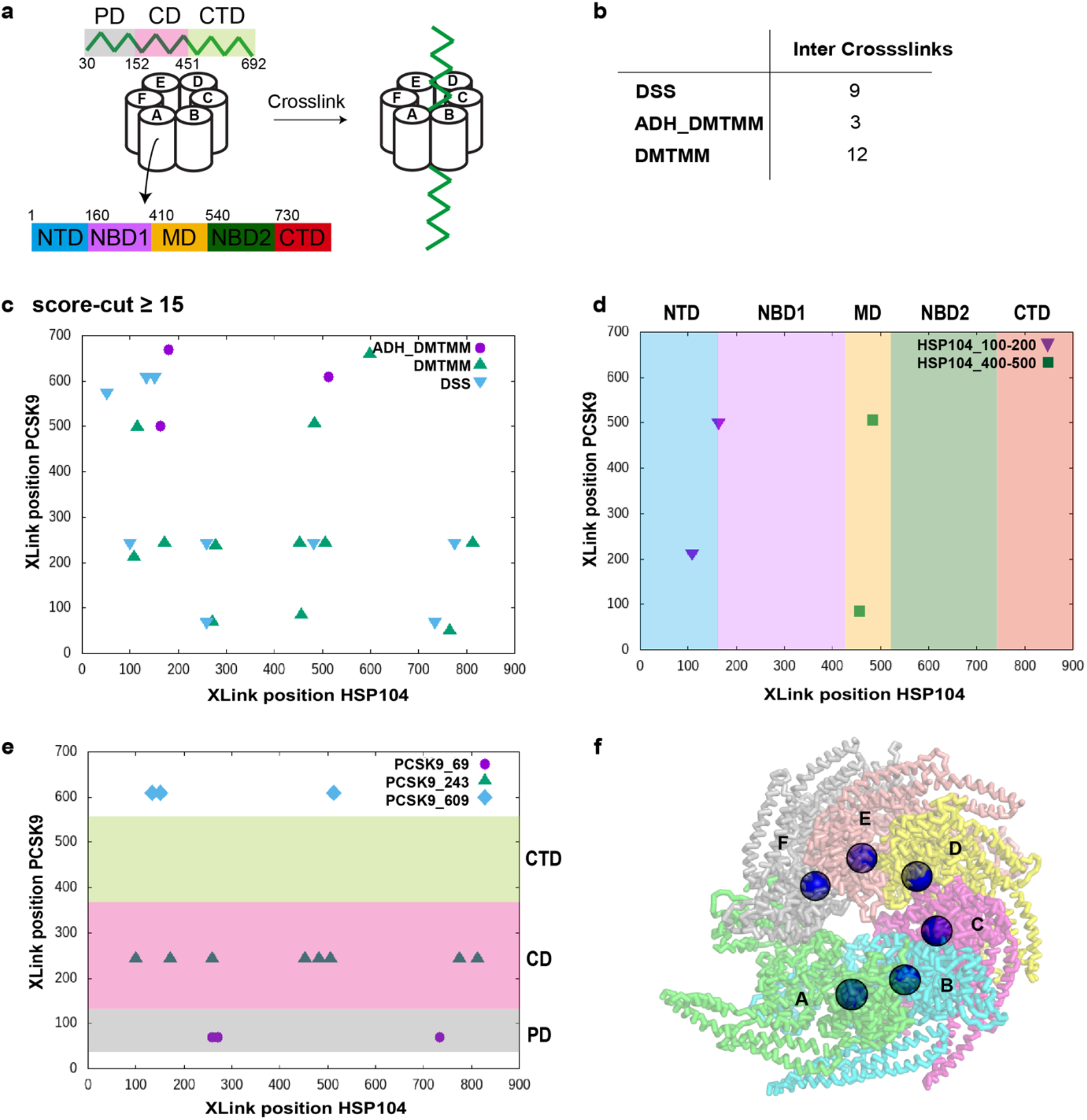
Hsp104 interacts with a substrate to reveal its disaggregating activity. **a**. Hsp104 was cross-linked with PCSK9 using commercially available crosslinking agents to determine its protein regions that interact with a substrate since Hsp104 is an ATP-dependent molecular machine that facilitates the disaggregates refolding. To reveal which region predominately binds the substrate, a schematic arrangement of Hsp104 and PCSK9 domains were presented in the Figure, where CD stands for catalytic domain, CTD – C-terminal domain, MD – middle domain, NBD1 and NBD2 – nucleotide-binding domains, NTD – N-terminal domain. It is believed that Hsp104 hexamer is active against aggregates which bind in the axial channel of the chaperone, hence it is essential to determine residues that crosslink to misfolded protein. **b**. Hsp104 was crosslinked with DSS, ADH and DMTMM. As a result, 9 inter crosslinks were formed using DSS, 3 inter crosslinks were generated using the mixture of ADH and DMTMM, and 12 inter crosslinks were detected after crosslinking with DMTMM. In each case a score cut has been applied, which amounted to 15 A. **c**. All inter crosslinks formed between Hsp104 and PCSK9 are presented in the chart. In this case, positions of amino acids that form a crosslink are plotted against each other to reveal Hsp104 and PCSK9 regions that are involved in crosslinking formation. The data plotted vertically represent residues of Hsp104 that form the particular constraints, while those plotted horizontally correspond to PCSK9 amino acids. Each crosslinking agent was marked with its symbol: (i) ADH_DMTMM_ZL is a purple circle, (ii) DMTMM_ZL is a green triangle and (iii) DSS is an inverted blue triangle. **d**. In Hsp104 protein, the largest possible number of crosslinks was formed within its N-terminal region and middle domain. **e**. In PCSK9, the largest number of crosslinks was formed within catalytic domain, however each of the domains exhibited ability to interact with Hsp104. Especially, 243 residue is an active chain that forms 8 crosslinks with Hsp104 residues. This may suggest that PCSK9 is translocated through various Hsp104 residues, included those localized in centrally placed channel of the Hsp104 structure. **f**. Residue that confirms the binding of PCSK9 in axial channel of Hsp104.

This also suggests that initial encounter complex is mediated by N-terminus not the C-terminus. Additionally, we find that the N-terminus of PCSK9 is more frequently crosslinked to Hsp104. Combined with a higher frequency of crosslinks of the substrates of NTD of Hsp104 suggest that the entire complex is predominately formed. (Fig. 5b) We determined 4 and 9 inter-molecular crosslinks within NTD and MD domains, respectively. Similarly, in PCSK9 sequence, we determined 3 residues which interact with several Hsp104 positions (Fig. 5e): (i) 69, (ii) 243 and (iii) 609. Especially, 243 position was crosslinked to 8 different amino acids of Hsp104, which suggests that entire polypeptide has an ability to translocate through various chaperone surfaces, including these residues located in axial channel that provides its disaggregating activity (Fig. 5f). Our crosslink data on Hsp104 bound to a model substrate uncovered a robust approach to begin mapping how Hsp104 may interact with substrates to reveal general mechanisms of Hsp104 activity.

## Discussion

Hsp104 is a protein that plays a critical role in disaggregation of stress-induced protein assemblies. In this study we employed Hsp104wt and Hsp104mt that forms a hexamer to determine all possible structural differences in these two complexes and to reveal Hsp104wt interactions with its prospective substrate in order to elucidate which protein regions are involved in the formation of such construct. We determined structural differences between Hsp104wt and mt, revealing that Hsp104mt exhibits more expanded structure whereby this Hsp104 genetic variant is less prone to (susceptible to) action of crosslinking agents (Fig. 6a and 6b). As a consequence, for each chemistry we determined smaller number of constraints in Hsp104mt. Such structural asymmetry is especially discernible between A and F subunits – using our approach we determined a distinctive constraint (634-258) that only fits these two subunits in both X-ray and cryo-EM wild type Hsp104 hexamer. In addition, we revealed which Hsp104wt regions and PCSK9 amino-acid residues are predominantly involved in the formation of Hsp104:PCSK9 complex. Two Hsp104wt regions (100-200 and 400-500) and three PCSK9 (69, 243 and 609) residues mostly interact with each other, giving (establishing) the disaggregation machinery that employs ATP to remove monomer units from protein assemblies.

**Figure 6.**
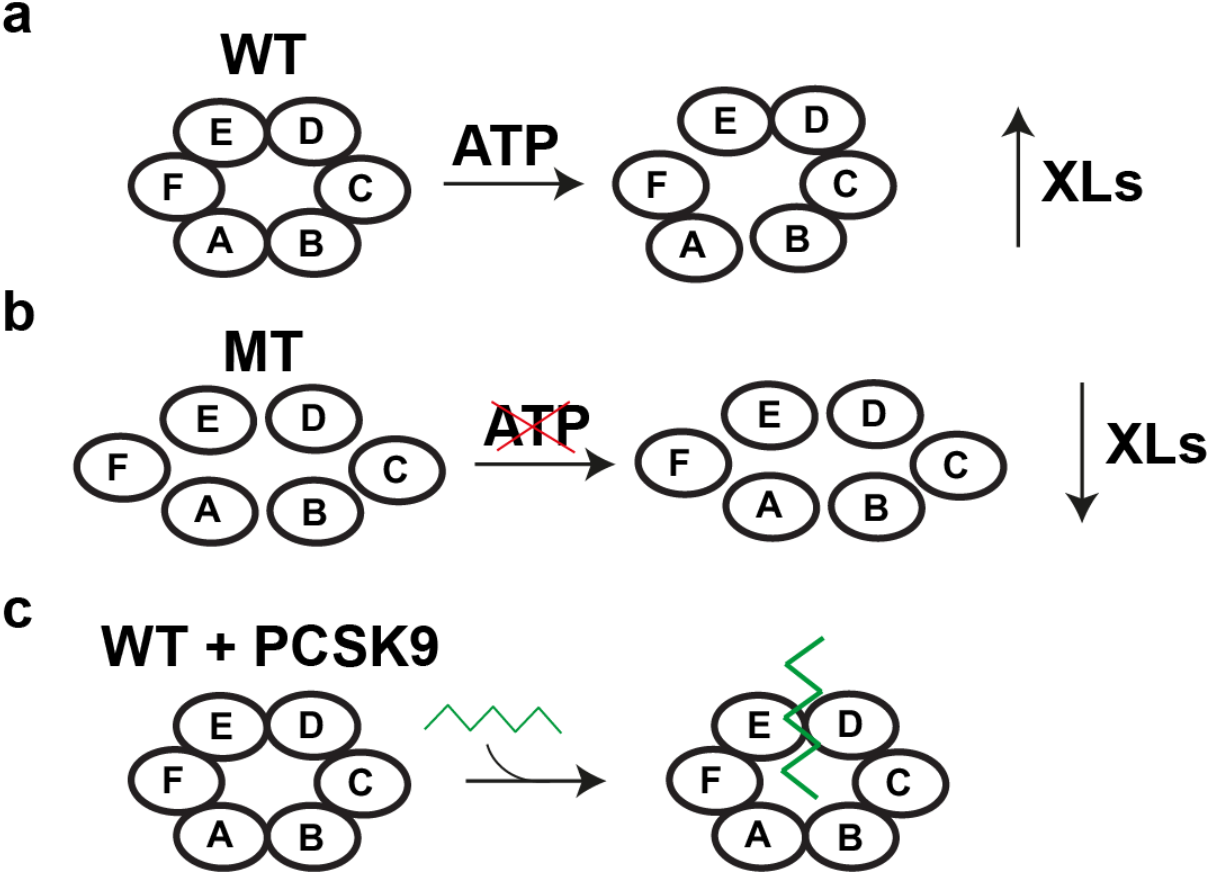
Asymmetry of Hsp104 detected by XL-MS. **a**. Wild type Hsp104 recognizes ATP which mediates disaggregation. This feature has an impact on the yield of crosslinking reaction. Moreover, Hsp104wt structure is more condensed. **b**. Genetically modified Hsp104 does not recognize ATP, which renders efficiency and yield of crosslinking reaction. Hsp104mt also has a looser structure and distances between particular units are larger, which inhibits the formation of cross-links. **c**. Hsp104wt interaction with PCSK9 yields the formation of several inter crosslinks, which indicates that Hsp104 recognizes PCSK9 as a substrate and translocates it via its axial channel.

### Different XL-MS chemistries are consistent with structures

It is commonly known that Hsp104 disassembles toxic aggregates of protein but also provides its protective function in several amyloid-related diseases. When expressed in yeast models, it promotes aggregation while its presence in animal cells may appear to regulate the clearance of amyloid assemblies. [32] In general, there are two main mechanisms of Hsp104 action. On the one hand, it uses its axial channel to disaggregate a substrate in an ATP-driven manner but also employs its NBD2 domain to capture entities before their aggregation. Such opposing and also cooperative mode of action requires to be revealed in easy, quick and affordable way in each study. [33,34] Our approach to determine any conformational changes or asymmetries as well as residues and protein regions that are involved in interaction seems to be especially useful in such experiments where particular agent may have an impact on Hsp104 activity and/or structure with a limitation that prior models of structures are essential for the interpretation. Here we employed two Hsp104 genetic variants (mt and wt) and structures (X-ray and cryo-EM) to reveal XL-MS application in geometric and functional studies. We determined that all the cross-links (generated using different chemistries) mapped onto both X-ray and cryo-EM structure fits perfectly to each of them – the only differences were caused by genetic variants of Hsp104 and thus Hsp104wt forms more links due to its more compact structure, while cross-linking reaction of Hsp104mt leads to the formation of lower number of constraints. In this case, we revealed that in such experiments it is possible to employ any X-ray or cryo-EM structure without losing any important data. Nowadays, there is conviction that X-ray crystallography and cryo-EM are two competitive structural techniques with differences such as sample preparation, accessibility and application, however one should keep in mind that structural determination is just a preamble to something bigger, which is combining structural information with biological functions.[35] Hence, if it is possible to gain similar data in our XL-MS approach using two different structures, there is no need to employ more expensive or less accessible technique.

### Visualization of substrates difficult using structural biology methods that rely on averaging but XL-MS uncovers discrete binding

Moreover, crosslinking reaction along with our novel analysis pipeline can be successfully employed in the elucidation of protein-protein interaction, especially when it is essential to precisely determine which regions are involved. Over the years, a lot of in-depth studies have aimed to determine how exactly Hsp104 hydrolyzes protein assemblies. [2,8,18,30,36,37] In general X-ray crystallography and cryo-EM are commonly employed in protein structure determination. After refinement, such an approach provides the average 3D model of protein and/or complex. [38,39] In our experiment, we visualized XL-MS data onto particular X-ray and cryo-EM structure thereby revealing discrete contacts that might be overlooked using commonly employed techniques. Having in mind that Hsp104 has an ability to alternate between two features (disaggregation and protection) [33] it is especially important to uncover each possible interaction of Hsp104 with its substrate, which helps to determine the exact model of Hsp104 activity.

### Conclusion

Our high-resolution XL-MS experiments combined with our analysis pipeline have uncovered exciting nuances to the dynamics of Hsp104 forming the staircase conformation only previously visualized with cryo-EM or X-ray. Furthermore, our experiments have uncovered robust methods to interpret substrate binding to gain insight into how substrates interact using XL-MS alone. Using XL-MS it is not only possible to confirm the protein structure but also to reveal its interaction with a substrate. Such an approach, based on the available cryo-EM or X-ray structure combined with XL-MS enables to determine all the residues that can bind a substrate. Future experiments can be focused on a cross comparison of substrates to identify general features of substrate binding and mechanisms of translocation.

## Acknowledgements

L.A.J was supported by a scholarship from the Effie Marie Cain Scholar in Medical Research. We would like to thank Andrew Lemoff from the UTSW Proteomics Core for his valuable insights on troubleshooting procedures in crosslinking mass spectrometry. We thank Dr. Andrzej Joachimiak (Argonne National Laboratories) for providing wt Hsp104, mt Hsp104 and Hsp104:PCSK9 complexes for XL-MS analysis. We thank all members of the Joachimiak lab, in particular Bryan Ryder, for discussions and input on the manuscript.

## Contributions

K.W. and L.A.J. initiated the project. K.W. performed all crosslinking experiments and analysis. K.W. and L.A.J. conceived of and directed the research and wrote the manuscript. All authors contributed to the revisions of the manuscript.

## Data availability

Raw XLMS data is available as source data 1. Any other data generated during and/or analyzed during the current study are available from the corresponding author on reasonable request.

## Materials and Methods

### Hsp104 expression and purification

*Calcarisporiella thermophila* wt Hsp104 (wt ctHsp104), ATP-binding deficient Hsp104 encoding a proline to alanine substitution at 383 (mt ctHsp104) and PCSK9 were kindly gifted by Dr. A. Joachimiak (Argonne National Laboratories). Hsp104 expression was induced at an OD 600 of 0.4–0.6 with 1 mM IPTG for 16 h at 15°C. Cells were harvested by centrifugation (4,000 g, 4°C, 20 min), resuspended in lysis buffer (40 mM HEPES-KOH pH 7.4, 500 mM KCl, 20 mM MgCl2, 2.5% glycerol, 20 mM imidazole, 2 mM β-ME) supplemented with 5 μM pepstatin A and complete protease inhibitor tablets (Roche). All purification steps were carried out at 4°C. Cells were lysed by incubation with 20 mg/mL hen egg lysozyme and sonication. Lysate was clarified by centrifugation at 16,000 rpm for 20 min and loaded onto Ni-NTA resin. The resin was washed with 10 volumes of wash buffer (40 mM HEPES-KOH pH 7.4, 500 mM KCl, 20 mM MgCl2, 2.5% glycerol, 20 mM imidazole, 2 mM β-ME). Protein was eluted in wash buffer supplemented with 350 mM imidazole. TEV protease was added to eluted protein, and the sample was dialyzed against wash buffer containing no imidazole for 4 h at room temperature followed by ∼16 h at 4°C. After dialysis and cleavage, the protein was loaded onto a second Ni-NTA column to remove the 6xHis tag and uncleaved protein. Eluted wt and mt Hsp104 was pooled, concentrated, and exchanged into storage buffer (20 mM HEPES-KOH pH 7.5, 200 mM KCl, 20 mM MgCl2, 10% glycerol, 2 mM ATP, and 1 mM DTT). A portion was used immediately for biochemical assays, and the remainder was flash cooled in liquid nitrogen and stored at −80°C until further use. PCSK9 was expressed and purified according to previously published protocol. [40]

### XL-MS of Hsp104 wt/mt and Hsp104 with PCSK9

Hsp104 wt/mt (0.3 mg/ml in 100 µl total volume) and Hsp104 with PCSK9 were crosslinked in dedicated buffers as follows:

|  | cross-linking agent | final concentration | shaking speed [rpm] | time [min.] | temperature [°C] | Quenching |
| --- | --- | --- | --- | --- | --- | --- |
| Hsp104 wt/mt | DSS | 1 mM |  | 1 | 37 | 50 mM AB at 37 °C for 30 min |
|  | DMTMM | 12 mg/ml | 750 | 15 | 37 | 200 mM AB for 30 min |
|  | ADH/ DMTMM | 8.3 /12 mg/ml | 750 | 15 | 37 | 200 mM AB for 30 min |
| Hsp104: PCSK9 + ATP | DSS | 1 mM |  | 1 | 37 | 50 mM AB at 37 °C for 30 min |
|  | DMTMM | 12 mg/ml | 750 | 15 | 37 | 200 mM AB for 30 min |
|  | ADH/ DMTMM | 8.3 /12 mg/ml | 750 | 15 | 37 | 200 mM AB for 30 min |

All samples separated elecrophoretically by SDS-PAGE (NUPAGE™, 4 to 12%, Bis–Tris, 1.5 mm or home-made SDS-Gel) and visualized by coomassie stain. After quenching, samples were lyophilized and resuspended in 8 M urea, which was followed by reduction with 2.5 mM TCEP (37 °C, 30 min) and alkylation with 5 mM iodoacetamide (RT, 30 min in darkness). Then the samples were diluted to 1 M urea with 50 mM (NH_4_)HCO_3_ (AB) and trypsinized (1:50 m/m, Promega) at 37 °C (overnight, 600 rpm). 2% (v/v) formic acid was added to decrease the pH of the solutions. Samples were then purified by solid-phase extraction (Sep-Pak tC18 cartridges, Waters®) and fractionated by size exclusion chromatography (SEC) using Superdex Peptide column. The collected fractions were freeze-dried and resuspended in water/acetonitrile/formic acid (95:5:0.1, v/v/v) to a final concentration of approximately 0.5 µg/µl. 2 µl of each sample was injected into Eksigent 1D-NanoLC-Ultra HPLC system coupled to a Thermo Orbitrap Fusion Tribrid system at the UTSW Proteomics core.

The analysis of the mass spectrum data was done by xQuest. Each raw data was first converted to open.mzXML format using mscovert (proteowizard.sourceforge.net). Search parameters were set differently based on the crosslink reagent. For DSS crosslink search: maximum number of missed cleavages (excluding the cross-linking site) = 2, peptide length = 5–50 aa, fixed modifications = carbamidomethyl-Cys (mass shift = 57.021460 Da), mass shift of the light crosslinker = 138.068080 Da, mass shift of mono-links = 156.078644 and 155.096428 Da, MS1 tolerance = 10 ppm, MS2 tolerance = 0.2 Da for common ions and 0.3 Da for cross-link ions, search in ion-tag mode. For DMTMM zero-length crosslink search: maximum number of missed cleavages = 2, peptide length = 5–50 residues, fixed modifications = carbamidomethyl-Cys (mass shift = 57.02146 Da), mass shift of crosslinker = − 18.010595 Da, no monolink mass specified, MS1 tolerance = 15 ppm, and MS2 tolerance = 0.2 Da for common ions and 0.3 Da for cross-link ions; search in enumeration mode. For ADH, maximum number of missed cleavages (excluding the crosslinking site) = 2, peptide length = 5–50 residues, fixed modifications = carbamidomethyl-Cys (mass shift = 57.021460 Da), mass shift of the light crosslinker = 138.09055 Da, mass shift of monolinks = 156.10111 Da, MS1 tolerance = 15 ppm, MS2 tolerance = 0.2 Da for common ions and 0.3 Da for cross-link ions, search in ion-tag mode. FDRs were estimated by xprophet [41] to be 0.1-0.46 and Cα-Cα distances were calculated using xwalk.

### Interpretation of crosslinks on experimental structures of Hsp104

Since each cross-link could be formed within one subunit or between two neighbouring subunits, we employed heat maps to order the distance values and determine which cross-links (intra, inter forward or inter reverse – for detail see Figure 2) were acceptable to be mapped on the particular structure. For Hsp104:PCSK9 complex only residues that formed inter cross-links were mapped onto Hsp104 structure to determine which protein region predominantly interacts with a substrate. All the crosslink mappings were visualized using Pymol, while the figures were prepared using custom Gnuplot scripts and Prism.

