## Supplementary Information for "Mapping Hsp104 structure and substrate interactions using crosslinking mass spectrometry"

**Source data 1. Raw XL-MS files for WT Hsp104, mutant Hsp104 and  WT Hsp104 in complex with PCSK9 substrate.**

### Supplementary Figures and Legends

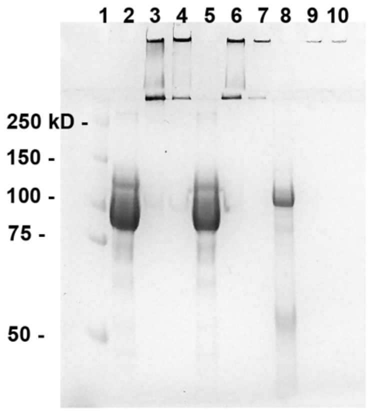


1 Protein ladder
2 Hsp104wt
3 Hsp104wt ADH_DMTMM
4 Hsp104wt DMTMM
5 Hsp104mt
6 Hsp104mt ADH_DMTMM
7 Hsp104mt DMTMM
8 Hsp104:PCSK9
9 Hsp104:PCSK9 ADH_DMTMM
10 Hsp104:PCSK9 DMTMM

**Supplementary Figure 1. SDS-PAGE of crosslinker HSP104 complexes.**
Hsp104wt/mt (i) as well as Hsp104 with its substrate (ii) were crosslinked in dedicated buffers ((i) – without ATP, (ii) supplemented with ATP) using ADH/DMTMM as crosslinking agent. Lines: 1 is a protein ladder (Precision Plus Protein Unstained Standard 10-250 kD, Bio-Rad), 2, 5 and 8 represent proteins before crosslinking reaction, while lines 3,4,6,7 and 9,10 correspond to crosslinked proteins.

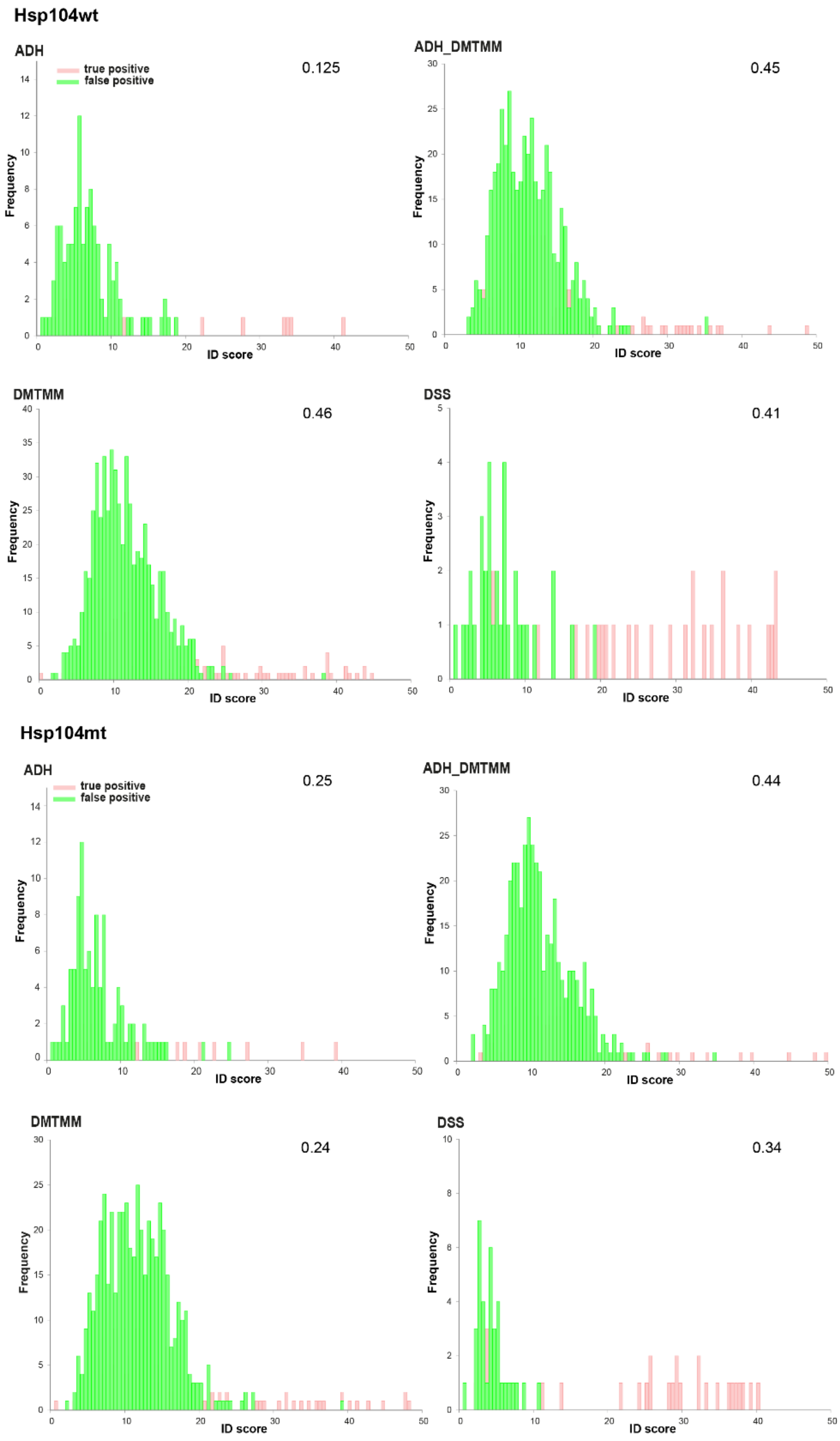


**Supplementary Figure 2. Wt Hsp104 and mt Hsp104 false discovery rate estimation.** Representative true positive (red) and false positive (green) distribution plots separated by score-id to calculate false discovery rates (FDRs) for each XL-MS dataset.

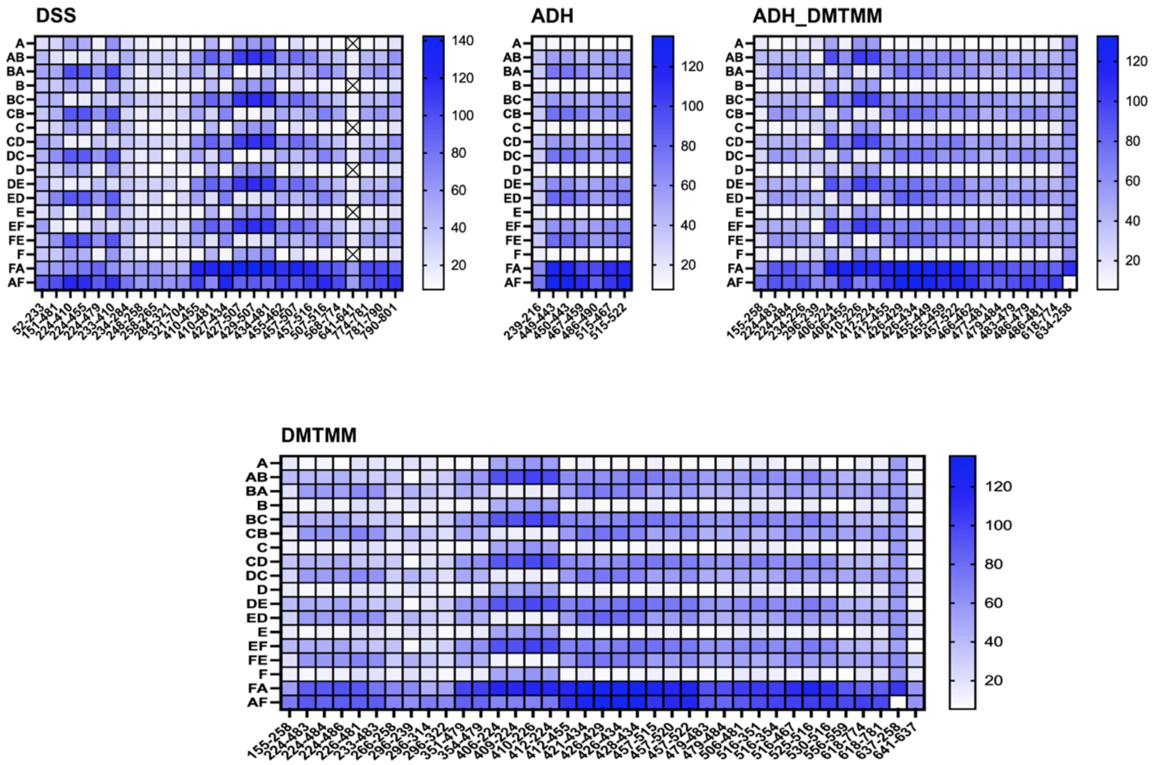


**Supplementary Figure 3. Mapping WT Hsp104 crosslinks onto ctHsp104 cryo-EM structure.** Heat maps showing cross-links formed within one unit or between two neighboring units in Hsp104 wt cryo-EM structure, using DSS, ADH_DMTMM and DMTMM. Coloring represents the distance between two linked residues, where the greater the distance, the darker the color.

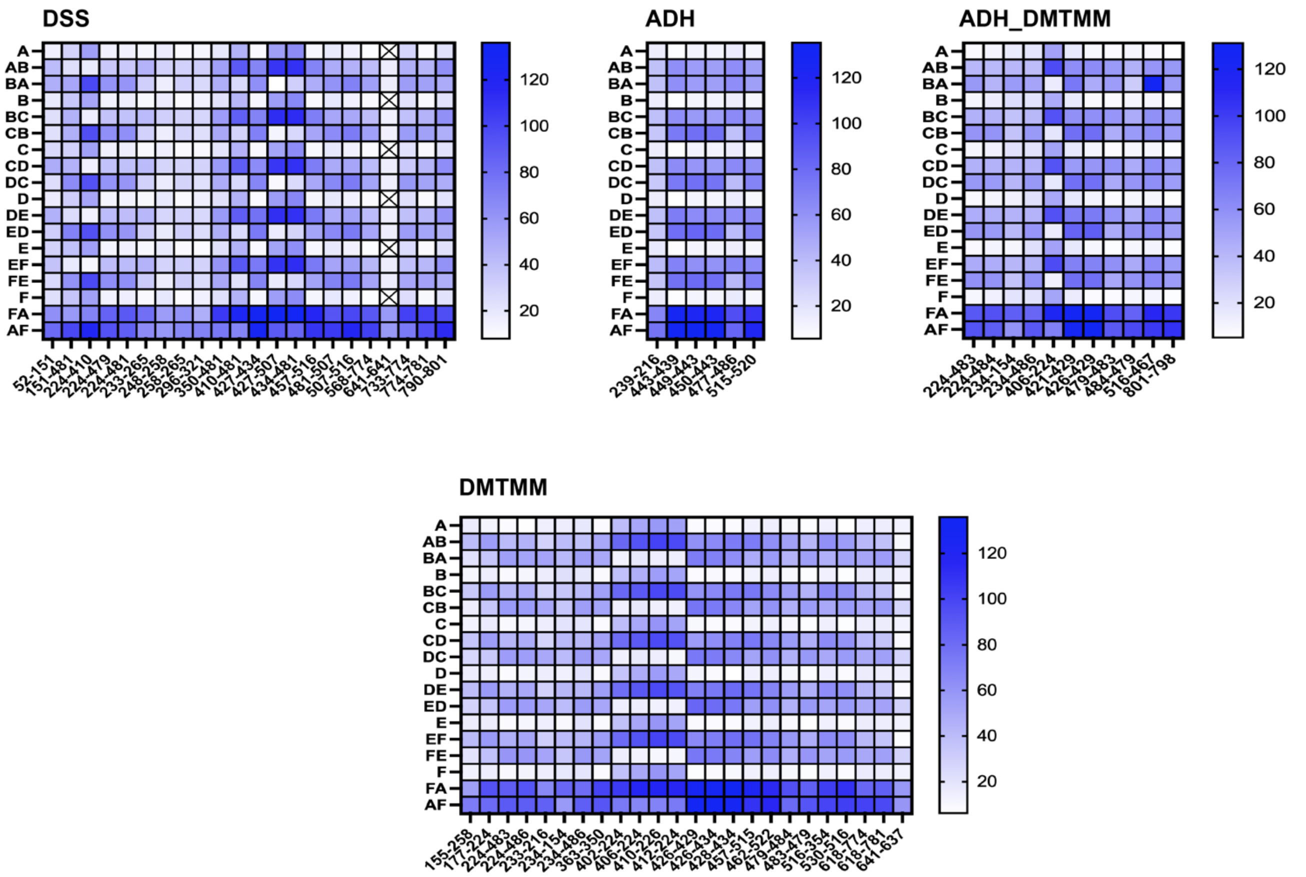


**Supplementary Figure 4. Mapping Mutant Hsp104 crosslinks onto ctHsp104 cryo-EM structure.** Heat maps showing cross-links formed within one unit or between two neighboring units in Hsp104mt cryo-EM structure, using DSS, ADH_DMTMM and DMTMM. Coloring represents the distance between two linked residues, where the greater the distance, the darker the color.

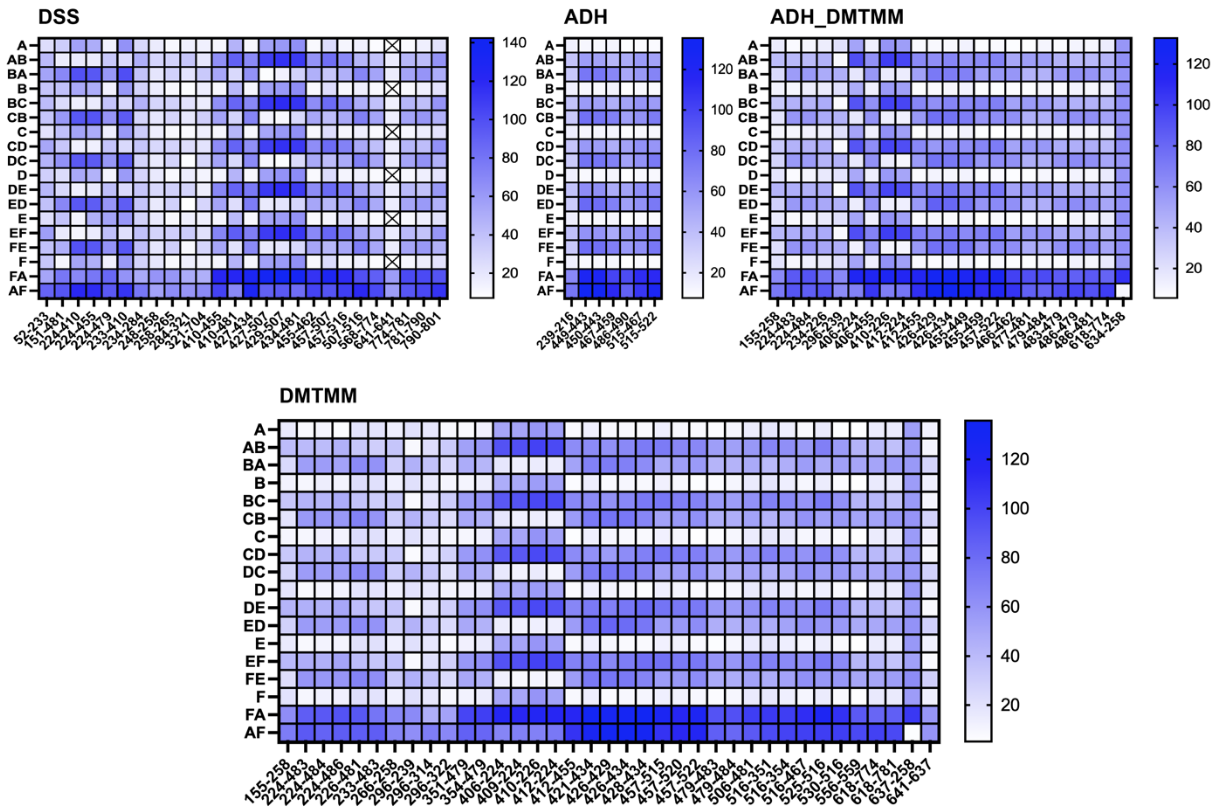


**Supplementary Figure 5. Mapping WT Hsp104 crosslinks onto ctHsp104 X-ray structure.** Heat maps showing cross-links formed within one unit or between two neighboring units in Hsp104 wt X-ray structure, using DSS, ADH_DMTMM and DMTMM. Coloring represents the distance between two linked residues, where the greater the distance, the darker the color.

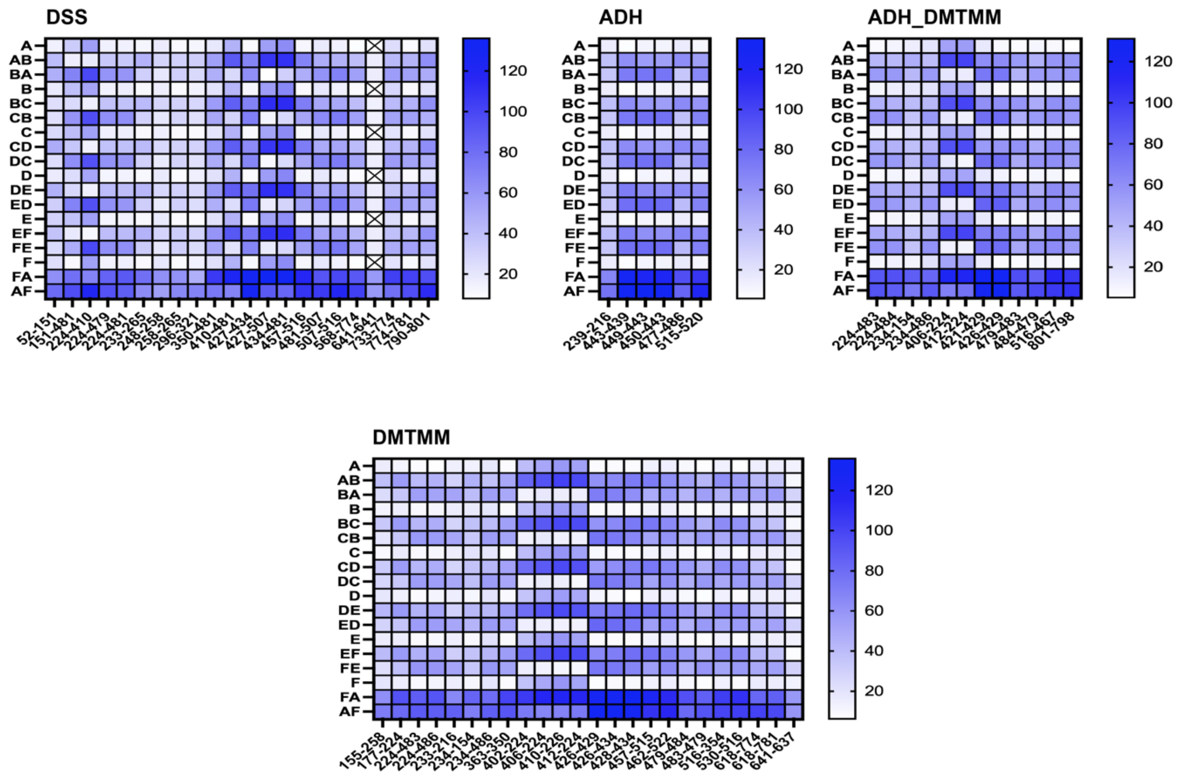


**Supplementary Figure 6. Mapping MT Hsp104 crosslinks onto ctHsp104 X-ray structure.** Heat maps showing cross-links formed within one unit or between two neighboring units in Hsp104mt cryo-EM structure, using DSS, ADH_DMTMM and DMTMM. Coloring represents the distance between two linked residues, where the greater the distance, the darker the color.

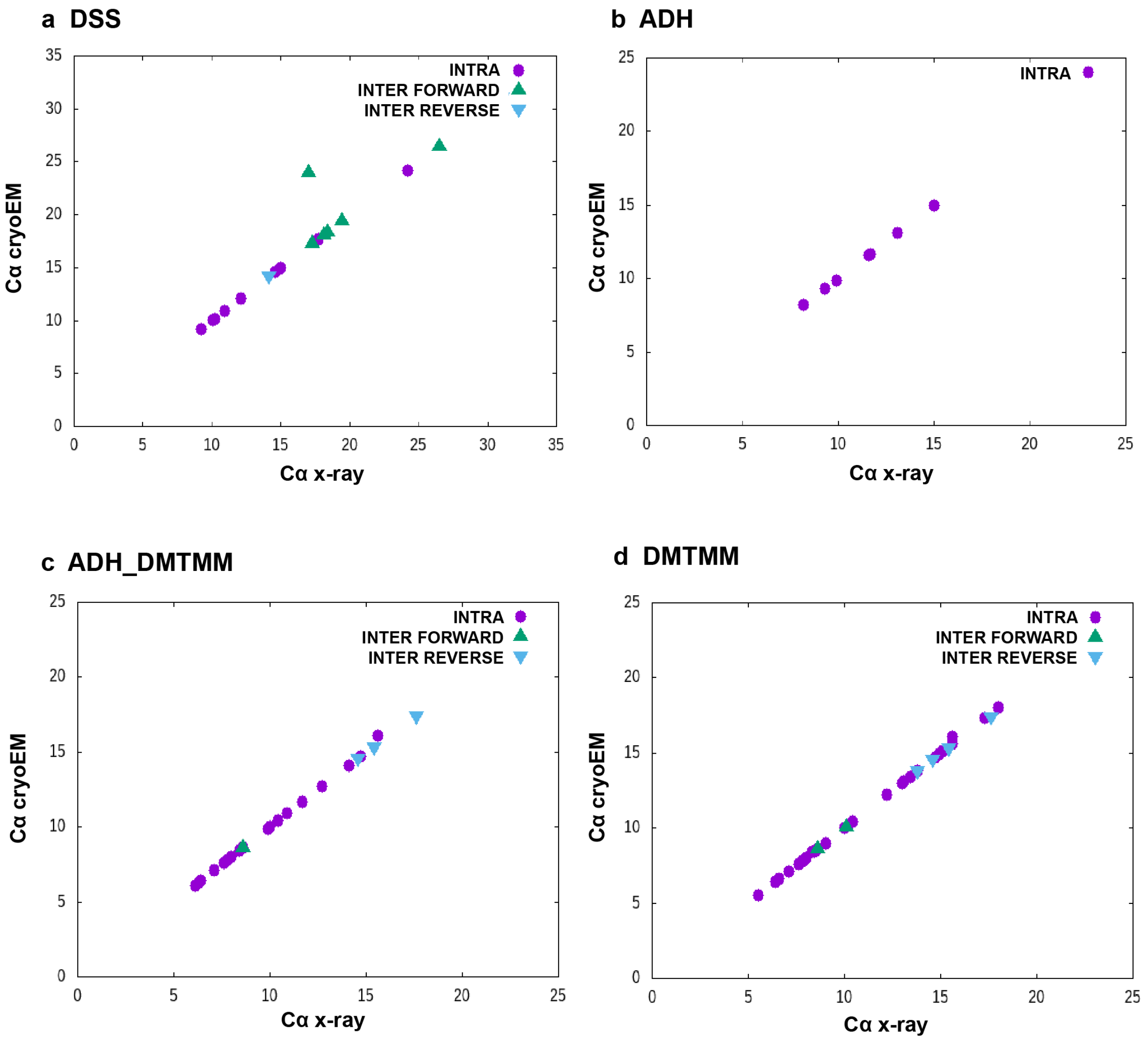


**Supplementary Figure 7. Agreement between calculated crosslink distances from WT ctHsp104 between the ctHsp104 X-ray and cryo-EM structures.**

Each cross-link distances mapped onto X-ray and cryo-EM structure were compared. Such an approach revealed that within crosslinking data obtained for Hsp104wt there is only one crosslink which differs from the other crosslinks (see green triangle in DSS plot). All the other crosslinks fit both structures.

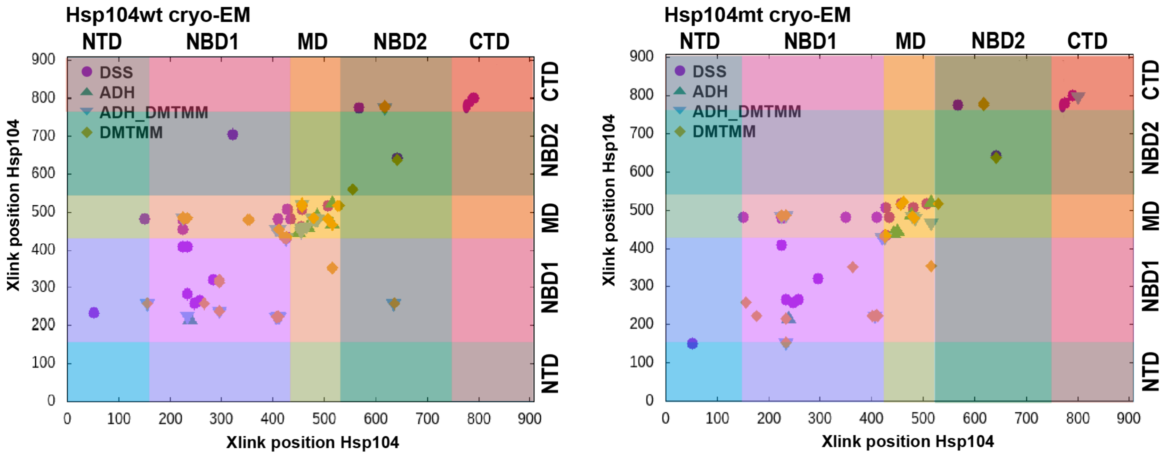


**Supplementary Figure 8. Hsp104 regions that predominantly form crosslinks.**
All crosslink positions were plotted against each other to reveal Hsp104wt/mt regions that are involved in the formation of constraints. In each case, the highest number of crosslinks was formed within NBD1 and MD domains. Only minor differences can be detected between genetic variations of Hsp104 (Hsp104mt forms fewer constraints), while crosslink distribution mapped onto particular structures does not reveal significant differences, which confirms structural similarity of Hsp104 cryo-EM and X-ray structure.

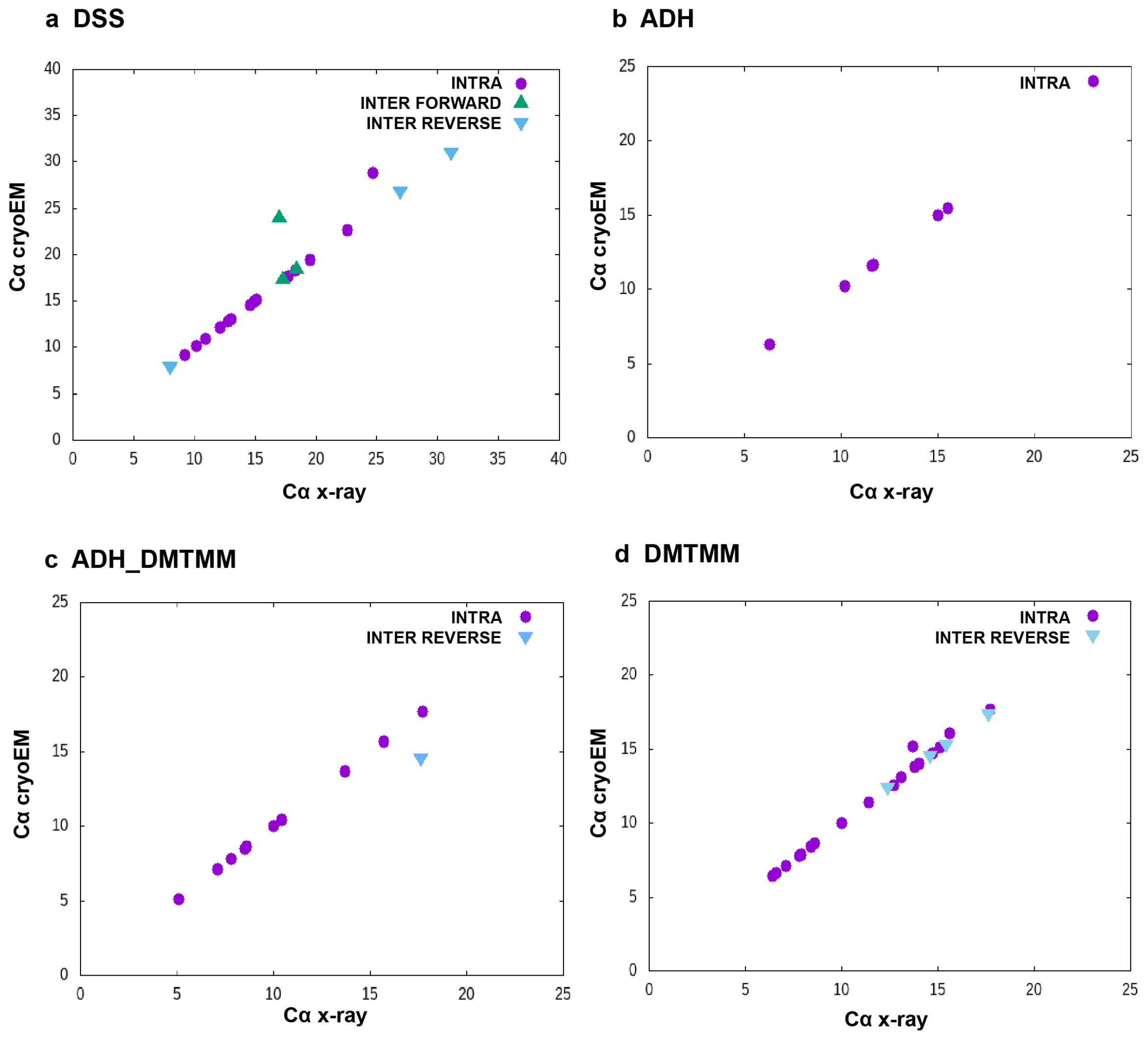


**Supplementary Figure 9. Agreement between calculated crosslink distances from MT ctHsp104 between the ctHsp104 X-ray and cryo-EM structures.**
Each cross-link distances mapped onto X-ray and cryo-EM structure were compared. Such an approach revealed that within crosslinking data obtained for Hsp104mt there are four crosslinks which differ from the others (see green triangle and purple circle in DSS plot, blue triangle in ADH_DMTMM_ZL plot and purple circle in DMTMM_ZL plot). All the other crosslinks fit both structures.

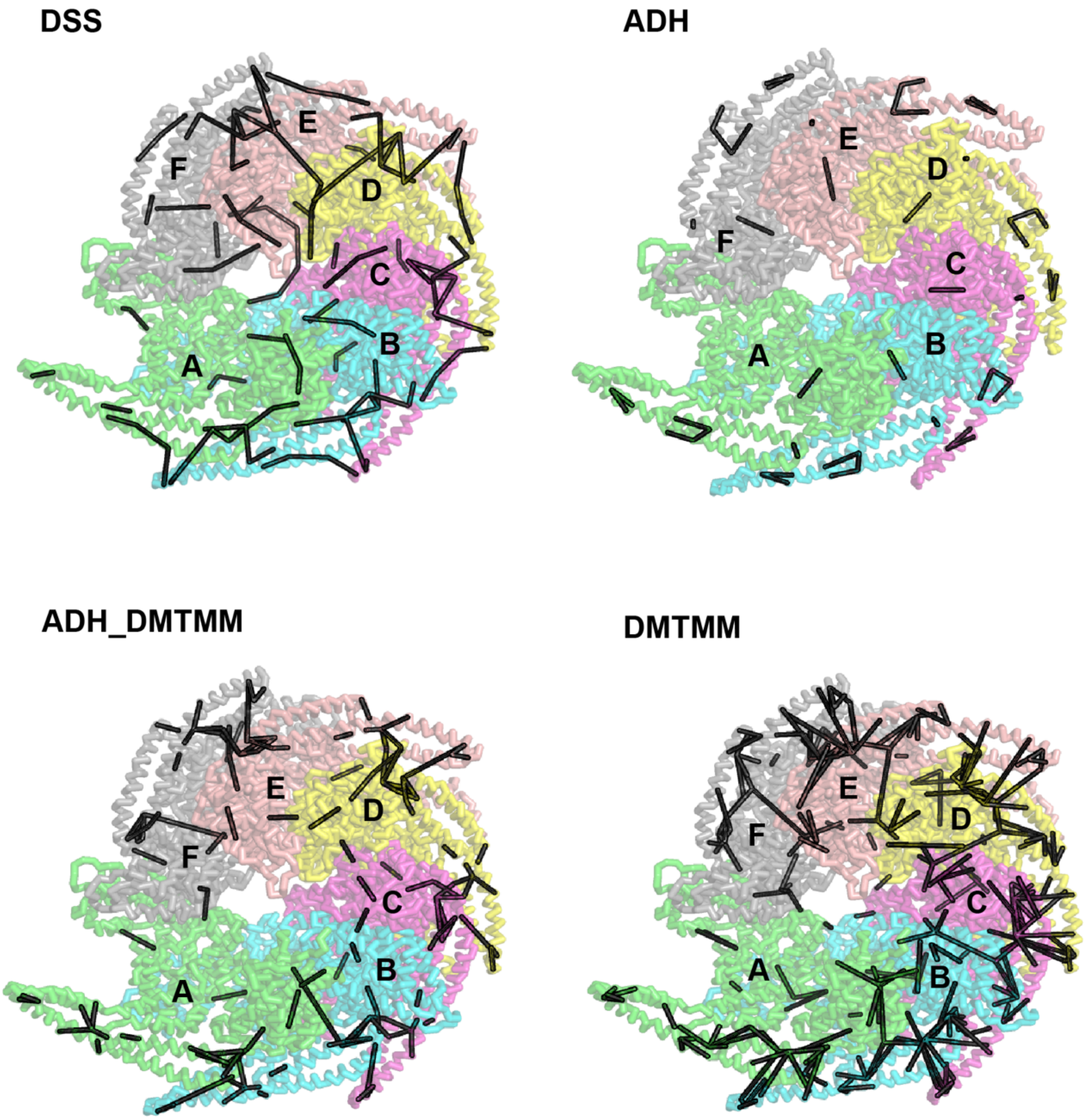


**Supplementary Figure 10. Mapping geometry-confirmed crosslinks WT ctHsp104 dataset onto the ctHsp104 cryo-EM structure.**
Representative of crosslinks mapped onto Hsp104wt cryo-EM structure. All the acceptable crosslinks were mapped onto cryo-EM structure to reveal protein geometry and determine which protein regions are predominantly involved in the crosslinking reaction. Since dataset obtained for both structure (cryo-EM and X-ray) and genetic variants (wild type and mutant) are similar, only mapping onto ctHsp104 cryo-EM structure is included.
